# Telomere Length as an Indicator of Lifestyle-Related Biological Aging in Hypertensive Adults: A Pilot Exploratory Study

**DOI:** 10.64898/2026.06.10.731327

**Authors:** Umair Aziz, Anum Zia, Anila Jaleel, Kiran Namoos, Sadia Farrukh, Saeeda Baig, Azra Mahmood

## Abstract

**Background:** Multiple epidemiological studies have given a global perspective that hypertension leads to changes in telomere length (TL). This study aimed to investigate the association between leukocyte telomere length and lifestyle factors among individuals with hypertension, a family history of the disease and healthy controls.

**Methods:** This pilot exploratory cross-sectional study included 45 participants (n = 15 per group) divided into hypertensive, familial hypertensive, and control groups. Lifestyle behaviors were assessed using the FANTASTIC Lifestyle Checklist, evaluating domains such as physical activity, diet, sleep, and stress. Relative telomere length (T/S ratio) of leucocytes was measured from peripheral blood samples using quantitative real-time PCR (qPCR). Data were analyzed using SPSS software, with ANOVA, Kruskal-Wallis test, and multivariable linear regression applied for statistical analysis.

**Results:** Relative telomere length differed significantly among the study groups (p = 0.008), with hypertensive participants demonstrating the highest median telomere-to-single-copy gene (T/S) ratio (2.66), followed by familial hypertensive (1.25) and control participants (0.87). Total lifestyle scores also varied significantly across groups (p < 0.001), with hypertensive individuals exhibiting higher mean scores (77.07 ± 5.59) than familial hypertensive (67.20 ± 3.00) and control participants (67.07 ± 5.97). A modest positive correlation was observed between total lifestyle score and relative telomere length (r = 0.329, p < 0.05), suggesting that healthier lifestyle behaviors in diagnosed hypertensive subjects following healthy lifestyles and regular medications were associated with longer telomeres. However, in multivariable linear regression analyses, lifestyle score, age, gender, hypertension status, and family history of hypertension were not independently associated with telomere length (all p > 0.05).

**Conclusion:** Lifestyle scores were positively associated with telomeres, and significantly higher telomere ratios in hypertensive subjects suggest complex biological interactions that require further investigation through larger longitudinal studies in the target population.

## Introduction

Hypertension is a highly prevalent cardiovascular risk factor among more than one billion people worldwide [1]. According to the World Health Organization (WHO, 2023), about 1.28 billion adults aged 30-79 years worldwide had hypertension, of which two-thirds live in low-and middle-income countries. Out of them, 46% are unaware of disease incidence [2]. According to the National Health Survey in Pakistan (NHSP), about 18.9% of people over 15 years of age are suffering from hypertension [3]. It is a polygenic disorder, which is not only genetic but is also attributable to environmental factors, i.e. gene and environmental interaction contribute to hypertension. Primary hypertension is due to an unknown cause and affects 90% of the hypertensive population. Secondary hypertension occurs secondary to an underlying organic disease including renal disease, adrenal gland disorders, pituitary diseases, and thyroid gland disorders. Hypertension, if left untreated, can lead to complications such as renal failure, myocardial infarction, retinopathy, and stroke [4].

Telomeres are the non-coding ends of linear chromosomes that protect the ends of the chromosomes from nucleases and prevent any loss of chromosomes or fusion of chromosomes, providing shelter or capping to the chromosomes. Telomeres consist of tandem repeating hexameric units of AGGGTT-rich base pairs. In autosomal cells with replication, the length of the telomere decreases after the completion of each cell cycle, resulting in the uncapping of chromosomes. In leukocytes, telomere length (TL) is longer at the time of birth, which decreases with age due to lifestyle modifications in adults. This decrease in telomere length represents the senescence of cells or the aging of the cells. [5]

National surveys suggest a correlation between leukocyte telomere length (LTL) and hypertension. Analysis of data from approximately 6,000 participants in the National Health and Nutrition Examination Survey (NHANES) revealed a non-linear relationship, with shorter telomere length being associated with an increased prevalence of hypertension [6]. Similarly, Zhang et al. reported that hypertensive patients who showed telomere lengthening during a 2.2-year follow-up had greater decreases in systolic blood pressure and pulse pressure in response to antihypertensive therapy [7]. These findings suggest that telomere dynamics may be influenced by blood pressure control over time.

Diet may also affect telomere biology. Liu et al. reported that in hypertensive older adults, telomere length and telomerase activity may be linked to hypertension, being influenced by environmental and pharmacological factors [8]. For instance, they observed that eating more vegetables may increase telomerase activity. Ultimately, combining clinical and lifestyle data often leads to mixed findings, likely due to variances in study design, target populations, and measurement techniques.

Research on this in Pakistan and South Asia remains quite limited. A case-control study by Rafiq et al. found that coronary artery disease patients in Pakistan had significantly shorter leukocyte telomeres, which may suggest that telomere attrition is linked to cardiovascular health in this population [9]. Familial factors appear to play a role as well. Farrukh et al. examined 204 father-newborn pairs in Pakistan, observing that fathers with hypertension and diabetes tended to have shorter telomeres, a trait that was also reflected in their newborns [10]. This points towards a potential paternal contribution to offspring telomere length. From a genetic perspective, a large-scale study of leukocyte telomere length by Delgado et al. involving 5,075 Bangladeshi adults identified a novel signal in the RTEL1 gene, while also confirming known telomere loci like TERC and TERT [11]. Together, these studies suggest that telomere dynamics are shaped by a complex mix of inherited and familial factors.

Lifestyle choices ranging from physical activity and stress levels to smoking and obesity are also believed to impact both blood pressure and telomere length. Supporting this, Nari et al. analyzed a Korean cohort of 9,474 people and found that a decline in a healthy lifestyle score was associated with a noticeable drop in overall quality of life [12].

Although several studies have linked unhealthy lifestyle behaviors to accelerated telomere shortening and hypertension, the evidence remains inconsistent, with some studies reporting weak or non-significant associations after adjustment for demographic and clinical factors. Furthermore, it remains unclear whether telomere shortening is a causal contributor to hypertension or primarily a consequence of chronic cardiovascular stress and aging. These uncertainties highlight the need for further investigation, particularly in underrepresented populations such as Pakistanis.

Therefore, the aim of the present pilot study was to compare leukocyte telomere length and health-related behaviors among hypertensive Pakistani individuals with and without a family history of hypertension and healthy controls. By examining the combined influence of familial predisposition and lifestyle factors on telomere length, this study sought to explore the potential relationship between telomere dynamics and hypertension in a Pakistani population.

## Materials and Methods

This pilot exploratory cross-sectional study included 45 participants (aged 25–60 years) divided into three equal groups: patients with hypertension and individuals with a family history of hypertension recruited from outpatient clinics at Shalamar hospital and healthy controls recruited from Shalimar hospital and Shalamar medical and dental college, Lahore between November 2024 and April 2025. The sample size was based on feasibility, availability of eligible participants, and study duration (pilot study). Participants were recruited by convenience sampling after receiving written informed consent. The study was conducted in accordance with the Declaration of Helsinki and approved by the institutional review board of Shalamar medical and dental college (IRB no. 0643). The participants were equally distributed into three groups (n = 15 /group). Group 1: individuals diagnosed with hypertension within 5 years, Group 2: normotensive individuals with a positive family history of hypertension (familial hypertensive) and Group 3: normotensive controls with no personal or family history of hypertension. Hypertension was defined according to the 2017 American College of Cardiology/American Heart Association guidelines as SBP ≥130 mmHg or DBP ≥80 mmHg on at least two occasions, or current use of antihypertensive medication [1]. Familial hypertensive participants had at least one first-degree relative diagnosed with hypertension but were themselves normotensive at recruitment. Individuals with endocrinological disorders such as: hyperthyroidism, cushing disease, acromegaly or pheochromocytoma, receiving any medications e.g. steroids, oral contraceptive pills, alcohol and pregnant or breastfeeding mothers were excluded. Potential confounders, including age, sex, body mass index (BMI), antihypertensive medication use, comorbidities, and lifestyle factors such as smoking, physical activity, and stress, were recorded and considered during interpretation.

A structured questionnaire was used to collect demographic and clinical data. Blood pressure was measured using a calibrated sphygmomanometer according to 2017 American College of Cardiology/American Heart Association guidelines. Lifestyle behaviors were assessed using the FANTASTIC Lifestyle Checklist, a validated multidimensional self-administered questionnaire that evaluates key lifestyle domains influencing health (https://cpb-cac1.wpmucdn.com/myriverside.sd43.bc.ca/dist/6/45/files/2014/01/Fantastic-Lifestyle-Checklist-Fillable-1smptgc.pdf). The checklist consists of items grouped under the following domains: Physical activity, Diet and nutrition, Smoking, Sleep, Stress, Social behavior and Career satisfaction. Each item was scored according to the standardized FANTASTIC scoring system, with higher scores reflecting healthier behaviors. Domain-specific scores were summed to generate a total lifestyle score, with possible scores ranging from poor to excellent lifestyle quality. Healthy lifestyle scores calculated from various modifiable risk factors have been used as a cost-effective, simple, and practical tool.

Genomic DNA was extracted using the Qiagen DNA mini kit method (Qiagen, Germany) from peripheral blood samples (5 ml) collected from all participants under aseptic conditions. Relative telomere length was measured using quantitative real-time PCR (qPCR), following the established telomere-to-single copy gene (T/S) ratio method [11]. The relative telomere length was calculated as the ratio of telomere repeat amplification to single gene amplification (T/S ratio), normalized to a reference DNA sample (for healthy controls). qPCR was performed using primers for telomeres and the single-copy gene, human beta-globin (HBG), as previously described by Cawthon (2009). The primer sequences were:

**Telomere Forward:** 5’-ACACTAAGGTTTGGGTTTGGGTTTGGGTTTGGGTTAGTGT-3’

**Telomere Reverse:** 5’-TGTTAGGTATCCCTATCCCTATCCCTATCCCTATCCCTAACA-3’

**β-globin Forward:** 5’-CGGCGGCGGGCGGCGCGGGCTGGGCGGCTTCATCCACGTTCACCTTG-3’

**β-globin Reverse:** 5’-GCCCGGCCCGCCGCGCCCGTCCCGCCGGAGGAGAAGTCTGCCGTT-3’

Amplification was performed on a real time PCR Thermocycler using EvaGreen qPCR Master Mix (Ref.: 2609-014/2025). Relative telomere length was calculated using the comparative C_t_ method, with the telomere target normalized against the reference gene.

The calibration method utilizes the Cawthon multiplex standard curve technique to calculate relative leukocyte telomere length. C_t_ handling splits threshold extractions by temperature, capturing the telomere signal at 74°C and the beta-globin signal at 88°C. Finally, quality control procedures mandate a spectrophotometric DNA purity ratio of ≥1.8 prior to qPCR, alongside on-plate monitoring using no-template controls and post-amplification melting curve analysis.

Possible sources of bias included selection bias due to hospital-based convenience sampling, recall bias from self-reported lifestyle data, and measurement variability in quantitative PCR-based telomere length assessment. Standardized procedures were used to minimize these biases. This study was conducted and reported in accordance with the STROBE guidelines for observational studies.

### Statistical Analysis

Statistical analyses were performed using SPSS software (version 25). For the data that followed a normal distribution, we used means and standard deviations, while we relied on medians and interquartile ranges (IQR) for non-normally distributed data. Categorical data were summarized using frequencies and percentages. To compare the three study groups, we used one-way ANOVA for the normally distributed variables and Kruskal-Wallis test for non-parametric data. Post-hoc pairwise analyses were carried out using Tukey’s Honestly Significant Difference (HSD) test, where applicable. For the categorical variables, we used either the chi-square or Fisher’s exact test, depending on the data distribution. Associations between relative telomere length and lifestyle scores were examined using Spearman’s rank correlation. Predictors of relative telomere length were evaluated using multivariable linear regression analysis. Statistical significance was defined as a p-value < 0.05.

Given the small sample size (n = 45) and multiple lifestyle domains examined, the analyses were exploratory. Accordingly, no correction for multiple testing was applied, and findings were interpreted with caution. Linear regression assumptions were verified, including normality and homoscedasticity of residuals. Multicollinearity was minimal, and model fit was assessed using adjusted R² values.

## Results

A total of 45 participants were included, divided equally (n=15 per group) into hypertensive, familial hypertensive, and control groups. While the median age differed among groups, with hypertensive participants being older (45 years) compared to familial hypertensive (40 years) and control participants (34 years); this difference did not reach statistical significance (p = 0.072).

There was no statistically significant difference in gender distribution across groups (p = 0.190); both hypertensive and familial hypertensive groups consisted of 9 males and 6 females, whereas the control group had 13 males and 2 females (Table 1).

**Table 1:** Demographic characteristics of the hypertensive, familial hypertensive and control groups.

| Variables |  | Hypertensive<br>(n=15)<br>Median (IQ) | Familial<br>Hypertensive<br>(n=15)<br>Median (IQ) | Control<br>(n=15)<br>Median (IQ) | p value |
| --- | --- | --- | --- | --- | --- |
| Age (years) |  | 45 (38–59) | 40 (34–49) | 34 (29–40) | 0.072 |
| Gender<br>n(%) | Male | 9(60.0%) | 9(60.0%) | 13(86.7%) | 0.190 |
|  | Female | 6(40.0%) | 6(40.0%) | 2(13.3%) |  |
| Blood<br>Pressure<br>mmHg | Systolic | 140 (130–150) | 120 (115–120) | 120 (115–120) | <0.001* |
|  | Diastolic | 90 (80–97.5) | 70 (70–80) | 80 (70–80) | <0.001* |
| Lifestyle<br>categories<br>n(%) | Fair | 0(0%) | 0(0%) | 1(6.7%) | 0.011* |
|  | Good | 10(66.7%) | 15(100.0%) | 14(93.3%) |  |
|  | Excellent | 5(33.33%) | 0(0%) | 0(0%) |  |
| Relative T/S ratio |  | 2.66 (0.95–4.14) | 1.25 (0.92–2.21) | 0.87 (0.64–1.52) | 0.008* |
| Total score of lifestyle<br>analysis (mean ± SD) | | 77.07 ± 5.59 | 67.20 ± 3.00 | 67.07 ± 5.97 | <0.001** |

Table 1 also showed that lifestyle category distribution differed significantly among the groups (p = 0.011). While most participants were categorized as having a “good” lifestyle, the hypertensive group included 5 participants with an “excellent” lifestyle and 10 with a “good” lifestyle. In contrast, the familial hypertensive group (15/15) and the control group (14/15) were predominantly classified in the “good” lifestyle category. Finally, relative telomere length expressed as the telomere-to-single gene (T/S) ratio differed significantly across groups (p = 0.008). The hypertensive group demonstrated the highest median T/S ratio [2.66], followed by the familial hypertensive group [1.25], while the control group exhibited the lowest values [0.87]. The total lifestyle score showed a significant difference across groups (p < 0.001).

Participants with hypertension demonstrated a higher mean lifestyle score (77.07 ± 5.59) compared to both the familial hypertensive group (67.20 ± 3.00) and controls (67.07 ± 5.97).

All 15 hypertensive patients enrolled in the study were receiving antihypertensive therapy. Among them, Calcium Channel Blockers (CCBs) were the most commonly prescribed class (40.0%), followed by ACE Inhibitors (ACEIs) (33.3%) and Angiotensin II Receptor Blockers (ARBs) (26.7%). The distribution of antihypertensive drug classes among the study participants is presented in Table 2.

**Table 2:** Distribution of antihypertensive drug classes among treated patients (n = 15)

| Antihypertensive Class | Number of Patients (n) | Percentage of Treated Patients (%) |
| --- | --- | --- |
| Calcium Channel Blockers (CCBs) | 6 | 40.00% |
| ACE Inhibitors (ACEIs) | 5 | 33.33% |
| Angiotensin II Receptor Blockers (ARBs) | 4 | 26.67% |
| Total | 15 | 100.00% |

Tukey’s HSD post hoc analysis (Fig 1) revealed that hypertensive participants showed significantly higher lifestyle scores compared with both familial hypertensive and control groups (*p* < 0.001), while no significant difference was observed between familial hypertensive and control groups.

**Fig 1.**
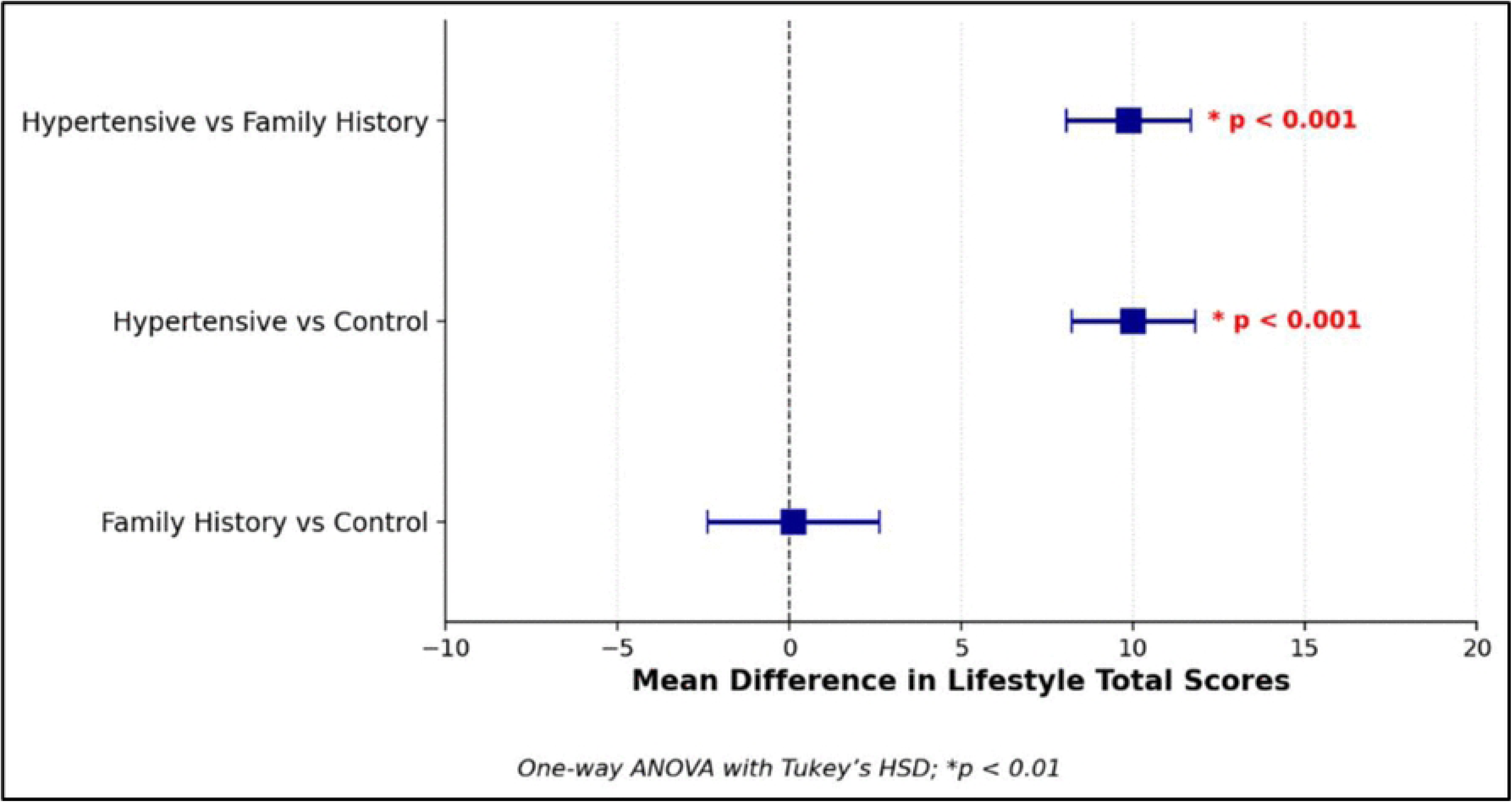
Pairwise comparison of lifestyle total scores among hypertensive patients, participants with a family history of hypertension, and healthy controls. The forest plot displays the mean differences in lifestyle scores between groups using one-way ANOVA with Tukey’s HSD post-hoc test. Data are presented as mean differences with 95% confidence intervals. Significant differences are observed between the hypertensive group and both the family history group and the control group (p < 0.001), while no significant difference is found between the family history and control groups.

The correlation matrix (Fig 2) showed a modest but significant positive association between relative telomere length and total lifestyle score (r = 0.329, p < 0.05), suggesting a better overall lifestyle is linked with longer telomeres. Total lifestyle score was strongly and positively correlated with physical activity (r=0.906), diet (r=0.872), sleep (r=0.941), stress levels (r=0.912), social behaviors (r=0.873), and job satisfaction (r = 0.924)( p < 0.001), while smoking showed a strong negative correlation (r=-0.78) with most healthy lifestyle domains. Age showed no meaningful correlation with telomere ratio and lifestyle score.

**Fig 2.**
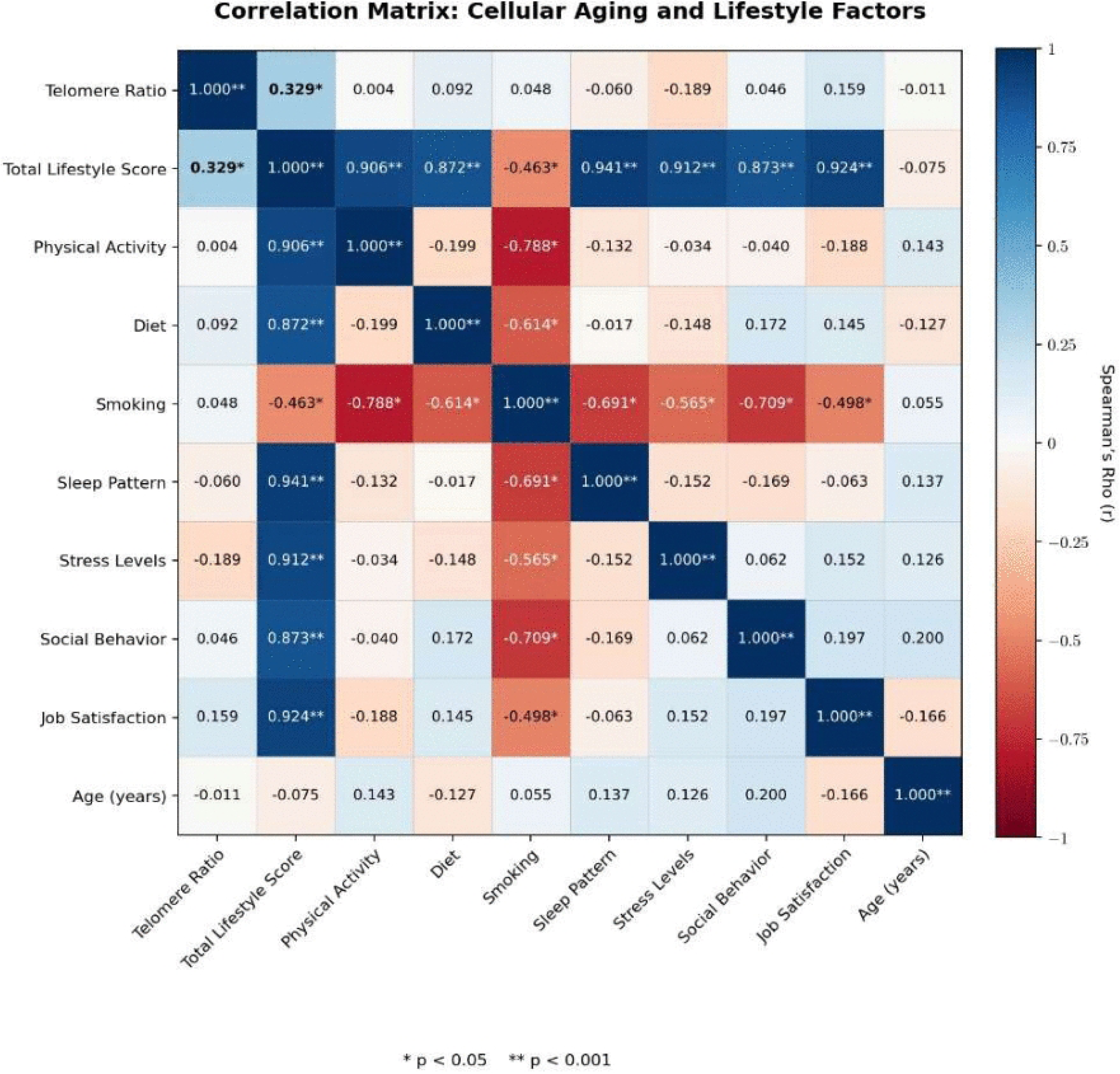
Spearman’s rank correlation matrix illustrating the relationships between telomere ratio, lifestyle factors, and age. The heatmap displays correlation coefficients (ρ) among the study variables. Color intensity indicates the strength and direction of the correlation, with blue representing positive and red representing negative associations. Asterisks denote statistical significance: *p < 0.05 and p < 0.001. A notable positive correlation is observed between the total lifestyle score and telomere ratio (r = 0.329, p < 0.05).

Table 3 presents the results of a multivariable linear regression analysis assessing predictors of relative telomere length. Total lifestyle score was not significantly associated with telomere length (β = 0.051, 95% CI: −0.065 to 0.168; p = 0.378), age (β = −0.005, p = 0.840), gender (β = 0.953, p = 0.138), hypertension status (β = 0.883, p = 0.340), and family history of hypertension (β = −0.626, p = 0.371) were also not significant predictors.

**Table 3:**
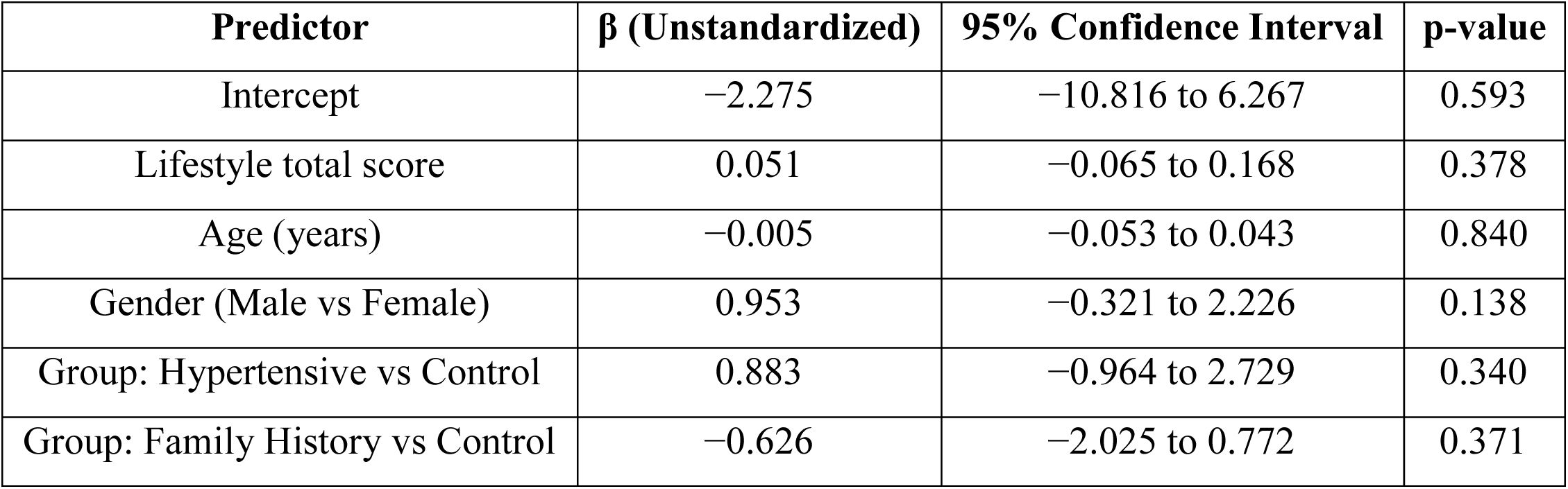
Multivariable linear regression analysis assessing predictors of relative telomere length.

A second regression model examining individual lifestyle components, including diet, sleep, stress, and career satisfaction, likewise showed no significant associations with relative telomere length (Table 4). None of the lifestyle domains, demographic variables, or group classifications independently predicted telomere length (p > 0.05)

**Table 4:** Regression model examining individual lifestyle components.

| Predictor | $\beta$ (Unstandardized) | 95% Confidence Interval | p-value |
| --- | --- | --- | --- |
| Intercept | -5.302 | -15.339 to 4.736 | 0.291 |
| Diet score | 0.097 | -1.556 to 1.750 | 0.906 |
| Sleep score | 1.036 | -0.800 to 2.872 | 0.260 |
| Stress score | -0.605 | -2.040 to 0.831 | 0.399 |
| Career satisfaction score | -0.267 | -1.904 to 1.370 | 0.742 |
| Age (years) | -0.004 | -0.051 to 0.043 | 0.858 |
| Gender (Male vs Female) | 0.912 | -0.345 to 2.170 | 0.152 |
| Group: Hypertensive vs Control | 0.801 | -1.019 to 2.621 | 0.379 |
| Group: Family History vs Control | -0.571 | -1.982 to 0.840 | 0.416 |

## Discussion

The study results demonstrated a complex interplay between lifestyle behaviors and clinical status in determining telomere length. A significant finding of this study is that hypertensive patients exhibited significantly higher median telomere ratios compared to the other two groups (familial hypertensive and healthy controls) (p = 0.008). This finding is unexpected and appears to contrast with broader epidemiological evidence suggesting an association between hypertension and accelerated cellular aging, often referred to as the “telomere paradox” [13]. However, rather than indicating a direct protective biological effect of hypertension, this finding may reflect post-diagnosis lifestyle modifications, pharmacological treatment, or variability inherent to the cross-sectional pilot design. Accordingly, no causal inference can be made from the present results. The hypertensive group also demonstrated significantly higher overall lifestyle scores (p < 0.001), particularly in the domains of diet, sleep, and physical activity. In the categorical analysis, five of fifteen hypertensive patients achieved an “Excellent” lifestyle score, whereas none in the other two groups fell into this category. This pattern further supports the possibility of behavioral changes following diagnosis, where patients may adopt healthier lifestyles in response to medical advice and regular follow-up.

Previous studies by Zhang et al. (2020) and Vasan et al. (2008) have reported that ARBs and ACE inhibitors may enhance telomerase activity, potentially contributing to telomere preservation or reduced attrition [7, 14]. Collectively, these observations raise the hypothesis that improved lifestyle behaviors, together with antihypertensive therapy, may create a more favorable cellular environment for telomere maintenance compared with untreated healthy individuals who do not undergo similar interventions. Nevertheless, these interpretations remain speculative and require confirmation in larger, longitudinal studies.

In the present study, all hypertensive participants were receiving antihypertensive therapy, predominantly Calcium Channel Blockers (CCBs), ACE inhibitors, and Angiotensin II Receptor Blockers (ARBs) (Table 2). Interestingly, hypertensive patients demonstrated comparatively longer telomere lengths, which may suggest a protective effect of antihypertensive treatment on cellular aging. Recent literature supports this observation, reporting that controlled hypertension under pharmacological therapy is associated with better telomere preservation compared to uncontrolled disease. The beneficial effects of antihypertensive agents may be mediated through reduction of oxidative stress, inflammation and endothelial dysfunction, particularly with CCBs and ARBs [15, 16]. Therefore, the relatively longer telomere length observed in our hypertensive cohort may reflect the beneficial impact of regular antihypertensive therapy and disease control, although the findings should be interpreted cautiously due to the small sample size and cross-sectional study design. However, evidence remains inconsistent, as some studies suggest that telomere shortening is more strongly influenced by cumulative cardiovascular risk and long-term disease burden rather than current treatment alone [17, 18].

In this study, no significant association was found between lifestyle total score and relative telomere length in the multivariable linear regression model (β = 0.051, 95% CI = 0.065 to 0.168, p = 0.378). Likewise, age, gender, and group status (hypertensive vs control and family history vs control) were not independent predictors of telomere length. Although unadjusted analyses suggested potential trends, these results indicate that lifestyle score is not an independent determinant of telomere length after controlling for confounders. This suggests that telomere biology may be influenced by more complex or unmeasured factors and may require larger samples to detect subtle effects.

Although lifestyle appeared to be a significant predictor in univariate analysis, it lost significance in multivariable models after adjustment for disease status. Importantly, the lifestyle tool used in this study reflects current, post-diagnosis behaviors rather than long-term exposures. In hypertensive individuals, these scores likely represent active disease management. Since telomere attrition occurs over decades, current lifestyle measures may not capture earlier exposures that shaped baseline telomere length. Overall lifestyle patterns over time may have broader biological effects, with cumulative exposure likely playing a more important role than isolated measurements. Supporting this, a recent study has linked unhealthy or declining lifestyle patterns with poorer quality of life, with additional effects of age, gender, and depression, particularly in persistently low lifestyle groups [12].

A significant difference in relative telomere length was observed among groups (p = 0.008). Notably, the hypertensive group (median age: 45 years) showed the highest telomere length (median T/S ratio: 2.66), whereas the younger control group (median age: 34 years) had shorter telomeres (0.87). This finding contrasts with most studies reporting shorter telomeres in hypertension and cardiovascular disease [12, 19]. However, emerging evidence suggests a more complex, possibly non-linear relationship between telomere length and hypertension. For example, Deng et al. (2022) reported that longer telomeres were associated with reduced cardiovascular risk but an increased risk of hypertension [20]. Similarly, Rubio-Carrasco et al. (2024) and Afsar et al. (2023) reported associations between longer telomeres and higher blood pressure [21, 22]. Other studies have also described non-linear relationships, where both very short and very long telomeres may be linked to adverse outcomes [23, 24].

Overall, several studies have reported associations between systolic blood pressure and telomere length (p = 0.039), with a borderline trend for diastolic pressure (p = 0.061). These findings raise questions about whether telomere length is influenced by blood pressure alone or by interacting metabolic and aging-related factors. Recent work has emphasized the importance of systolic blood pressure control in older adults, suggesting that even small changes in management may affect telomere dynamics and biological aging [21]. Similarly, studies in dyslipidemia have shown mixed effects on telomere length, with both increases and decreases observed depending on lipid status and patient characteristics [25, 26].

Overall, current evidence suggests that telomere length is influenced by a combination of genetic, clinical, and lifestyle factors rather than hypertension alone. Age, gender, and lifestyle may contribute to telomere dynamics but do not appear to independently determine telomere length [19]. Further longitudinal studies focusing on hypertension treatment and metabolic interventions are needed to better clarify the role of telomere biology in cardiovascular health.

Earlier studies have demonstrated that telomere length (TL) is influenced by genetic factors; however, environmental exposures, including lifestyle behaviors, aging, and other non-genetic risk factors, are major contributors to telomere attrition [11]. Environmental influences may modulate this association by affecting oxidative stress, inflammation, and cellular aging processes. Some studies suggest that telomeres may serve as a potential biological marker for cardiovascular conditions, particularly hypertension, during the early stages of physiological aging in middle age. The other important factors investigated related to both TL attrition and oxidative stress were dyslipidemia and insufficient sleep [25, 27, 28]. A study reported that higher body mass index (BMI) was associated with shorter LTL, and higher low-density lipoprotein cholesterol was associated with longer LTL [29]. Another study revealed that maintaining high cardiorespiratory fitness (CRF) may neutralize the influence of obesity and CVD risk factors on telomere length, potentially leading to telomere elongation [30]. Similarly study by Benfield et al., (2026) showed that significantly longer telomere were observed in black individuals compared to white, but hypertension was more prevalent in the black population despite this longer telomere length [31]. These studies are consistent with the findings of our study in an Asian population, in which hypertensive participants exhibited a higher relative telomere length (T/S ratio) compared with individuals with a family history of hypertension and healthy controls. Therefore, focusing on a single lifestyle factor is insufficient, and a holistic evaluation of multiple lifestyle determinants is required to better predict disease risk and support effective lifestyle interventions.

In our study gender distribution was comparable across groups, with males comprising 29.0% of both hypertensive and familial hypertensive groups, whereas females were 42.9% (p = 0.190). Overall, previous studies suggest that telomere length (TL) tends to decrease with advancing age; however, variations may exist between males and females, and TL may also be influenced by several factors, including the use of medications, particularly antihypertensive drugs. A study comparing sexes found that only in females there is a statistically significant relationship between TL and age (r = −0.170, p value = 0.0146) [21]. This study also points out towards a significant association between systolic blood pressure control and TL, and a trend for diastolic blood pressure, suggesting hypertension management impacts TL. The findings also suggested a stronger relationship between DBP control, age, and HDL-C levels in females compared to males. Furthermore, age appeared to have a greater impact on SBP control in females (p = 0.01) than in males (p = 0.080) [30].

Hypertension has become a major public health concern in Pakistan, with a rapidly increasing prevalence estimated at approximately 46.2% [32]. Rising prevalence of hypertension among younger generation with compromised compliance needs urgent attention. There is need to identify the risk factors of hypertension for adequate prevention. Our results, combined with previous studies, suggest that non-genetic factors like lifestyle score contribute to hypertension and early intervention can improve and boost overall vascular health and its influence on TL.

Future studies with larger sample sizes, prospective longitudinal designs, and detailed assessment of genetic, environmental, and treatment-related factors are needed to clarify the relationship between telomere length and hypertension. Further research in different South Asian populations may also help determine whether the patterns observed in this study are population-specific or representative of broader biological mechanisms underlying hypertension

## Conclusion

Hypertension is a multifactorial disease influenced by hereditary predisposition and environmental factors. In this study, family history was used to assess familial risk rather than direct genetic analysis. Hypertensive patients on regular treatment and healthier lifestyle modifications showed longer relative telomere lengths, although these factors did not independently predict telomere length. Moreover, normotensive individuals with a family history of hypertension did not differ significantly in telomere length from healthy controls. These findings suggest that both disease management and modifiable lifestyle factors may contribute to cellular aging processes. Future prospective longitudinal studies incorporating genetic analyses are needed to further clarify the relationship between hypertension and telomere dynamics.

This study has several limitations. The use of convenience sampling from outpatient clinics may have introduced selection bias. Hypertensive participants, who were already engaged with healthcare services, were more likely to have received lifestyle counseling and medical treatment. This health-seeking behavior likely explains their higher lifestyle scores compared with community-recruited controls, indicating that the groups may not reflect baseline lifestyle patterns.

Other limitations include differences in age, sex, body mass index (BMI), medication use, and lifestyle factors, as well as the relatively small sample size. Moreover, the study was restricted to a specific age group and geographic region, which may limit the generalizability of the findings.

## Acknowledgements

We are grateful to the administration of Shalamar Medical and Dental College for their support in conducting this study. We also sincerely thank the Director of Research and Innovation for providing access to laboratory facilities and research support.

During the preparation of this work, the authors used ChatGPT (OpenAI) to improve the readability and language of the manuscript. All content was reviewed and approved by the authors.

## Author contributions

Conceptualization: Anila Jaleel, Azra Mahmood, Saeeda Baig. Formal analysis: Kiran Namoos.

Investigation: Anila Jaleel, Umair Aziz, Anum Zia. Methodology: Saeeda Baig.

Project administration: Anila Jaleel.

Resources: Azra Mahmood, Umair Aziz, Anum Zia. Supervision: Anila Jaleel, Sadia Farrukh.

Validation: Kiran Namoos, Sadia Farrukh. Visualization: Saeeda Baig.

Writing - original draft: Anila Jaleel, Umair Aziz, Anum Zia.

Writing - review & editing: Azra Mahmood, Kiran Namoos, Sadia Farrukh.

